# Training deep learning models on personalized genomic sequences improves variant effect prediction

**DOI:** 10.1101/2024.10.15.618510

**Authors:** Adam Y. He, Nathan P. Palamuttam, Charles G. Danko

## Abstract

Sequence-to-function models have broad applications in interpreting the molecular impact of genetic variation, yet have been criticized for poor performance in this task. Here we show that training models on functional genomic data with matched personal genomes improves their performance at variant effect prediction. Variant effect representations are retained even when fine tuning models to unseen cellular contexts and experimental readouts. Our results have implications for interpreting trait-associated genetic variation.

## Main

Deep learning models have rapidly become the state of the art for predicting marks of regulatory function from genome sequence. Despite accurately predicting both the pattern of chromatin marks and gene expression across genomic loci, however, several recent studies have highlighted significant limitations in the ability of some models to predict differences between individuals [1, 2]. Existing models struggle most with gene expression predictions, due in part to challenges integrating the *cis*-regulatory effects of distal enhancers [3, 4]. Performance is better in predicting the impact of *cis*-regulatory variation on local features, including transcription initiation, chromatin accessibility, and activity in reporter assays [5–7]. However, performance of state-of-the-art models does not yet saturate accuracy even in local prediction problems.

Most sequence-to-function models are trained using a single haploid reference genome as input [4, 8–13]. While this approach is computationally tractable and enables the accumulation of large datasets where matched genome sequences are not known, training on a single genome prevents models from observing the impact of genetic variation on genome function, potentially resulting in poor performance in variant interpretation tasks. Consequently, the extent to which training models using personalized genomes improves performance has become a subject of intense interest [14–16].

We recently described CLIPNET [17], a deep convolutional neural network trained to use input DNA sequences to predict maps of transcription initiation measured using PRO-cap, an assay that maps the 5’-capped ends of nascent RNAs [18]. We trained CLIPNET using functional data and matched diploid genomes from 58 Yoruban lymphoblastoid cell lines (LCLs) (+9 biological replicates) [19]. In addition to accurately predicting transcription initiation at single-nucleotide resolution from genome sequences, CLIPNET can correctly impute the effects of genetic variants on both transcription initiation quantity (tiQTLs) and directionality (diQTLs) [19].

Here, we asked whether training CLIPNET using matched personal genome sequences improved its ability to predict the impact of genetic variation on transcription initiation. We first trained CLIPNET models on random subsets of the 67 PRO-cap libraries and their associated personal genomes used in our original study. When predicting across loci on holdout chromosomes, predictions of the profile (distribution of PRO-cap reads within a given prediction window) and the total quantity of transcription initiation rapidly improved in accuracy as the number of training individuals increased, with diminishing returns beginning at around 10-20 individuals (Fig. 1B). We observed a similar trend when assessing prediction accuracy on personalized genomic sequence near genetic variants associated with proximal *cis* changes in transcription initiation. CLIPNET required training on 20-30 individuals before saturating accuracy in predicting the average of individual-level PRO-cap signal near tiQTL and diQTL lead SNPs (Fig. 1C).

**Figure 1:**
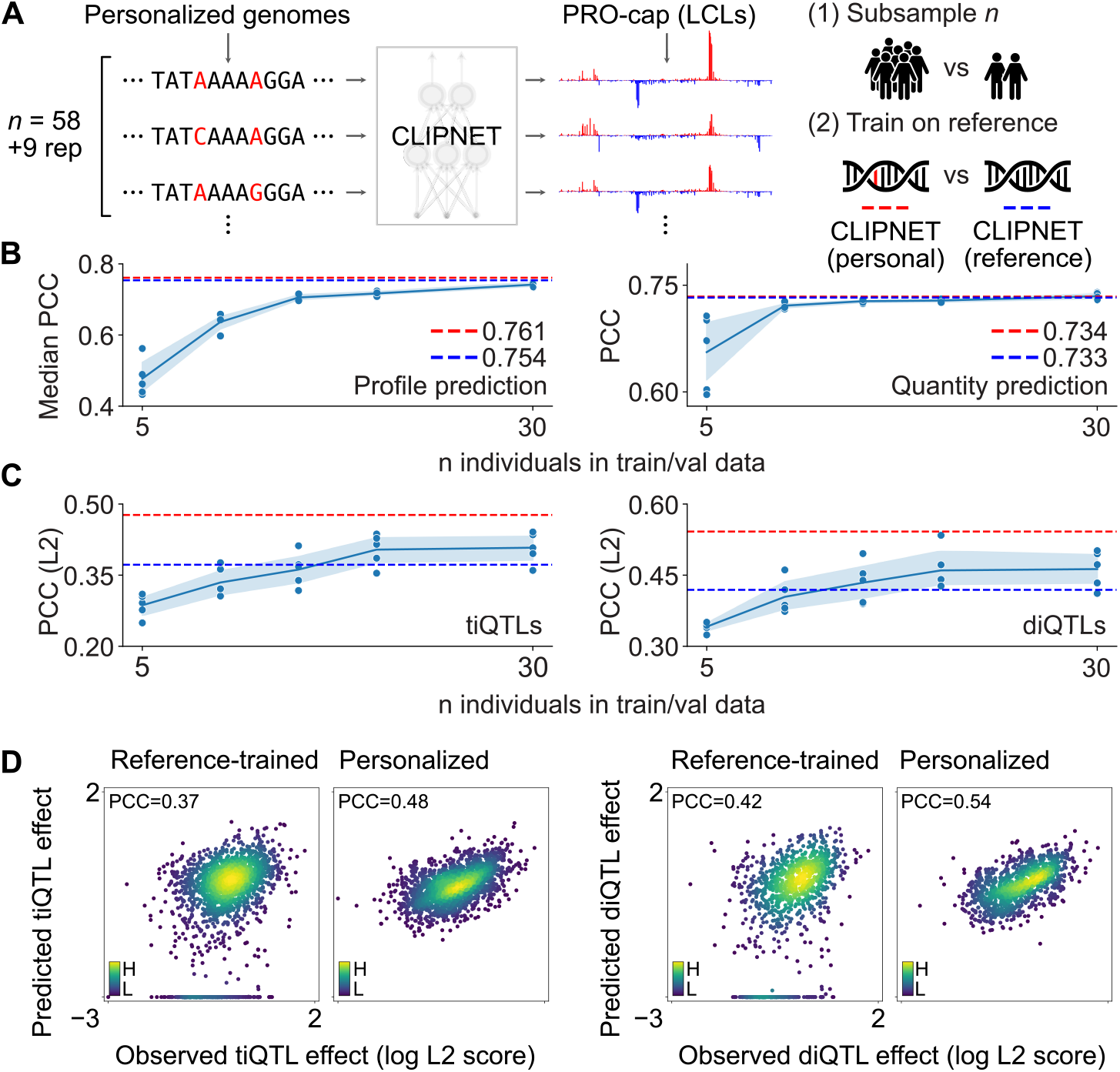
Training deep learning models on personalized genomic sequences improves variant effect prediction. (**A**) Schematic of the personalized genomes training scheme used for CLIPNET and the two ablation strategies (training data subsampling and variant masking/reference training). (**B**) Evaluation of subsampled models on transcription initiation profile (left) and quantity (right) prediction (*n* = 4598 loci). PCC = Pearson’s correlation coefficient. (**C**) Evaluation of subsampled models on predicting effects of tiQTLs (left, *n* = 2057) and diQTLs (right, *n* = 1207) (PCC between observed and predicted *L*^2^ variant effect scores). In both **B** and **C**, the performance of the original CLIPNET model is indicated by red dashed lines, while the performance of the reference-trained CLIPNET model is indicated by blue dashed lines. Individual subsampled model performances are indicated by dots, while the mean and 95% confidence intervals are illustrated by line plots. (**D**) Comparison of tiQTL (left two panels) and diQTL (right two panels) effect prediction by reference-trained (left panel in each pair) and personalized (right panel in each pair) CLIPNET models. Points are colored by a Gaussian kernel density estimate.

While the subsampling test indicated that training on more individuals improves model accuracy, these results could also be explained by CLIPNET benefiting from more training data, irrespective of whether the sequence inputs come from personalized genomes or reference genomes. To directly measure the contribution of training on personalized genomes to model performance, we trained a CLIPNET model on the full 67 library dataset [17], using only reference genome sequence with the genetic variants masked. The reference-trained CLIPNET model performed nearly identically to the personalized CLIPNET model and outperformed most of the subsampled models at predicting initiation profiles and quantities across loci in the genome (Fig. 1B, Supplementary Fig. S1). However, the reference-trained model substantially underperformed the full personalized CLIPNET model at predicting QTL effects, performing roughly on par with subsampled models trained on only 10 or 20 individuals (Fig. 1C-D). Notably, the reference-trained model predicted 17.0% (*n* = 350) of tested tiQTLs and 20.0% (*n* = 241) of tested diQTLs as having no effect (Fig. 1D). These results show that training on larger numbers of functional genomic datasets improves performance across loci, but performance on variant effect prediction tasks improves further by using dataset-matched personal genomes.

We next asked whether the performance gain from using larger functional datasets and matched personal genomes generalizes across cell types and experimental readouts. We used massively parallel reporter assays (MPRAs) to as a ground truth for regulatory SNP impact. Although PRO-cap and MPRAs measure distinct biological processes, transcription initiation and enhancer activity are correlated [20, 21], suggesting that models predicting SNP effects on initiation should also perform reasonably well in predicting enhancer activity.

We used fine tuning to adapt CLIPNET to a K562 PRO-cap dataset [22] and evaluated the accuracy of SNP effect prediction against a massively parallel reporter assay (MPRA) ground truth [23] (Fig. 2A). The transfer-learned CLIPNET K562 model showed strong correlations with experimental PRO-cap data, performing similarly to ProCapNet [10], a model trained natively on K562 PRO-cap data using the reference genome, in predictions across loci (Supplementary Fig. S2). Despite similar performance across loci, CLIPNET significantly outperformed ProCapNet in distinguishing expression-modulating variants (emVars) from non-emVars (Fig. 2B). Both PRO-cap-based models significantly outperformed Enformer [9], a large multi-task transformer-based model that has been criticized for poor performance on predicting personal gene expression [1, 2].

**Figure 2:**
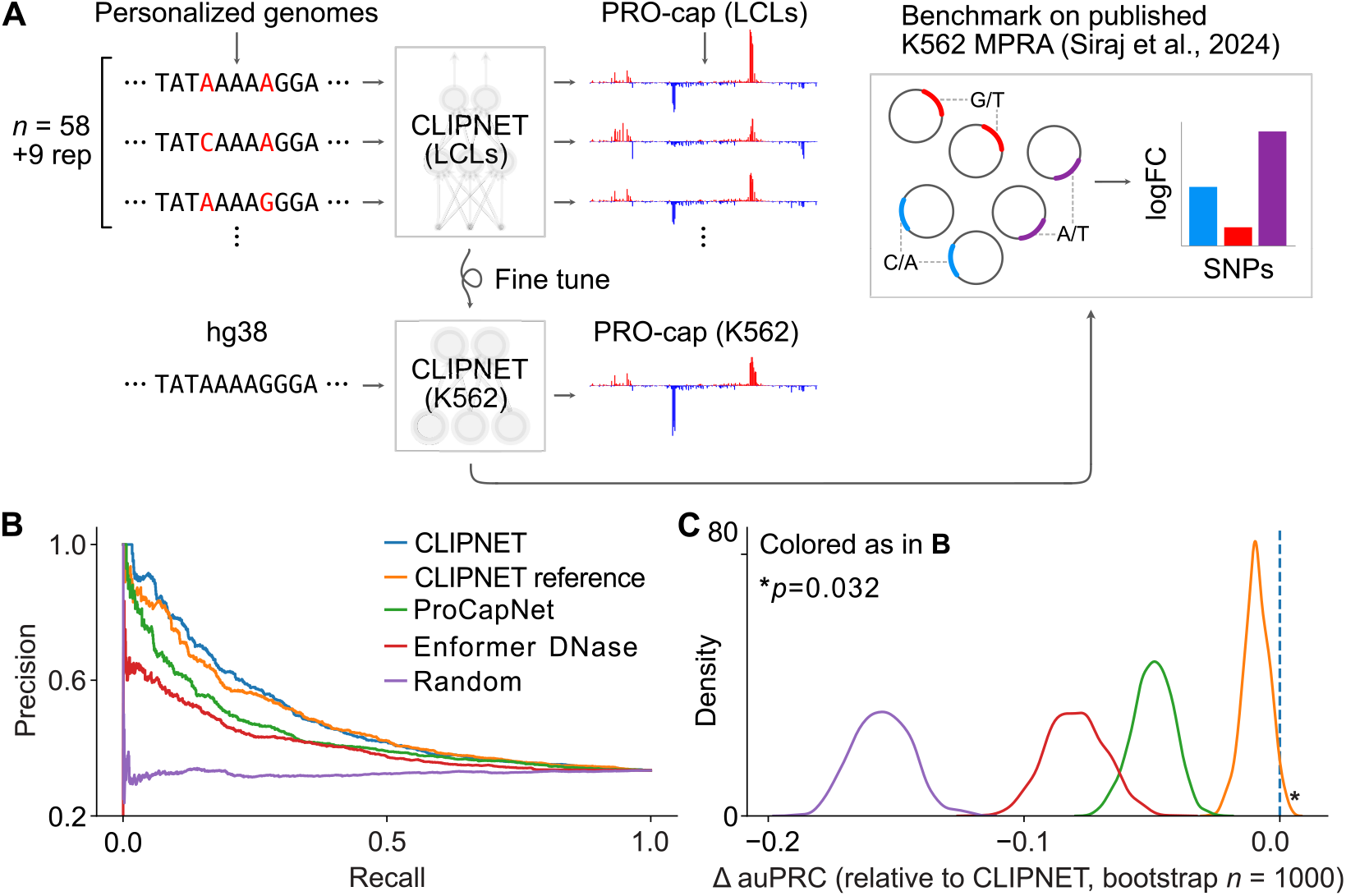
CLIPNET fine tuned to predict initiation in K562 outperforms models trained on only reference genome sequences at predicting MPRA variant effects [23]. (**A**) Schematic illustrating the procedure for fine tuning CLIPNET to predicting initiation in K562 and for benchmarking on MPRA variant effect prediction. (**B**) Precision-recall curves for all tested models on classifying expression-modulating variants (emVars, *n* = 2038) vs expressed non-emVars (*n* = 4057). To avoid data leakage from pretraining CLIPNET on personal genomes, only variants from test chromosomes 9, 13, 20, and 21 were analyzed. (**C**) Bootstrapped estimates of change in area under the precision-recall curve of all other models compared to CLIPNET.

To assess whether the variant representations learned by training on personalized genome sequences improved variant effect predictions, we also transferred the reference-trained CLIPNET model to K562. Again, the personalized CLIPNET K562 model outperformed the reference-trained CLIPNET K562 model at variant effect prediction, albeit by a smaller margin than was observed in the LCL initiation QTL prediction task (p = 0.038; bootstrap test; Fig. 2B-C). Next, we compared the change in transcription initiation of each SNP predicted by the model to the fold-change measured using the MPRA. These results reinforce our initiation QTL analysis and suggest that training on personalized genome sequences allows CLIPNET to observe substantial activity differences between nearly identical input DNA sequences, which reference-trained models fail to observe.

Our work has several valuable applications. First, MPRAs are frequently used to identify and validate candidate causal SNPs from genetic association studies [23–27]. Achieving reasonable *in silico* performance by training on both large functional genomic datasets and matched personal genomes has widespread potential for interpreting the molecular effects of trait-associated SNPs and prioritization for experimental validation. Second, enhancer design is an emerging field [28–31]. Models such as CLIPNET and ProCapNet, which predict transcription initiation from local sequences, are particularly well suited for designing and interpreting enhancer elements, which often show a stereotypical divergent initiation pattern in mammals [18, 21, 32, 33].

Recent work has shown that while fine-tuning gene expression predictions using genetic variation can improve predictions for individual genes, this performance uplift does not generalize to unobserved genes [15, 16]. Expression models are most sensitive to promoters and capture the effects of distal enhancers much less accurately [3, 34, 5]. These results make intuitive sense: learning accurate long-range models that capture the effect of distal enhancers requires fitting two complex and independent functions. First, the impact of DNA sequence on local *cis*-regulatory activity, and second, the complex logic by which different *cis*-regulatory elements integrate their signals across a locus. Both of these complex functions must be learned from a limited set of only 20,000 genes. Adding to the challenge, most of the highly expressed genes across different cell types—where mistakes are penalized heavily for models trained across different cell types or tissues—are housekeeping genes, which are less likely to be impacted by enhancers [35]. We argue that this complexity, combined with the limited set of training examples, makes the problem even more difficult.

Our results suggest a path forward for developing accurate variant interpretation models that predict gene expression. We propose that the first step is to accurately learn the effect of variants on local *cis*-regulatory activity. Our findings show that local models are substantially improved by incorporating genetic variation and by using larger datasets with multiple independent measurements of the same cell or tissue type. Genetic variation enhances the model’s ability to generalize to unseen variants and even to distinct experimental readouts, such as MPRAs. Accurate local models can then be integrated into models trained to predict gene expression, either through heuristics [5] or through long-range transformer-based models trained to learn regulatory logic. We also suggest that training more complex models may benefit from adding biological constraints, for instance by initializing model parameters using experimental chromatin contact data from assays such as Hi-C or Micro-C. This stepwise approach provides a more feasible path to addressing the inherent complexity of gene regulation models, laying the groundwork for better generalizability and numerous downstream applications interpreting trait-associated genetic variation.

## Methods

### LCL PRO-cap and personal genome processing

We downloaded and processed PRO-cap and personal genome data for 58 genetically distinct LCLs (+ 9 replicates) as described in the original CLIPNET manuscript [17]. Relevant components of the protocol are reproduced and summarized here.

We downloaded raw PRO-cap read counts from Gene Expression Omnibus accession GSE110638 [19] and phased genotypes from the 2019 1000 Genomes Project release (https://ftp.1000genomes.ebi.ac.uk/vol1/ftp/data_collections/1000_genomes_project/release/20190312_biallelic_SNV_and_INDEL/). To ensure consistency between the PRO-cap and genotyping data, we lifted over the PRO-cap libraries from their original hg19 reference to hg38.

We generated individualized genomes by applying the genotyped SNPs from each individual to the hg38 reference genome. Indels and structural variants were excluded due to their rarity and the complexity that they would introduce. To represent diploid genomic sequences, we used a “two-hot” encoding scheme; that is, we encoded each individual nucleotide at a given position using a one-hot encoding scheme, then represented the unphased diploid sequence as the sum of the two one-hot encoded nucleotides at each position. The sequence AYCR (= A(C/T)C(A/G)), for example, would be encoded as [[2, 0, 0, 0], [0, 1, 0, 1], [0, 2, 0, 0], [1, 0, 1, 0]].

PRO-cap peaks were individually called in each library using a pre-publication version of PINTS [32] supplied by the authors. As *cis*-regulatory elements commonly consist of two divergently transcribed core promoters spaced roughly 110 bp apart [18], we filtered for unidirectional PRO-cap peaks on each strand that were no more than 200 bp away from a unidirectional peak on the opposite strand. We then extracted 1kb, randomly jittered by up to 250 bp, of matched genomic sequence and PRO-cap tracks around the center of each PINTS call.

To enable model ensembling, we partitioned the genome along chromosomal boundaries into 10 roughly equally sized folds. We set aside fold 0 (consisting of chromosomes 9, 13, 20, and 21) for final evaluation of the model ensemble. The remaining 9 folds were then used to train 9 replicate models, each of which used a distinct holdout fold. This ensures that prediction quality at each position within the genome can be fairly evaluated using individual models.

### Subsampled model training

We randomly selected *n* = 5, 10, 15, 20, 30 PRO-cap libraries from the 67 used to train CLIPNET, then trained subsampled CLIPNET models on matched personal genome sequences and PRO-cap tracks (obtained as above) following the methods described in our previous manuscript (detailed in the following paragraph) [17]. We avoided sampling multiple isogenic replicates from the same individual in each subsampling experiment. For each *n*, we performed 5 such runs, resulting in a total of 25 subsampled CLIPNET models.

Each subsampled CLIPNET model consists of an ensemble of 9 replicate convolutional neural networks (CNNs) trained in a leave-one-out fashion using the 9 training folds described above. Individual CNNs use the CLIPNET architecture [17], which was inspired by the BPNet [8] architecture. The main body of the network consists of two convolutional layers (64 filters, width 8 and 128 filters, width 4), followed by a tower of 9 exponentially dilated convolutional layers (64 filters, width 3, dilation factors from 1 to 512) separated by skip connections. Batch normalization was applied after each convolutional layer. Rectified linear activations (ReLU) were used for each convolutional layer except for the first, which utilized an exponential linear activation. Max pooling (width 2) was applied after each of the first two convolutional layers and after the dilated convolution tower.

We applied two output heads after the convolutional tower. One head (profile) predicts base-resolution distribution of PRO-cap reads and consists of a single fully-connected layer, while the other (quantity) predicts the total quantity of PRO-cap reads in the output window and consists of an average pooling layer followed by a fully-connected layer. Batch normalization and ReLU activation were applied after each fully-connected layer. To jointly evaluate the prediction accuracies of these two output heads, we used a multiscale loss function.

Specifically, for a given 500 bp output window, let 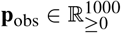 represent the base-resolution PRO-cap coverage and 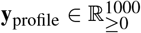 and 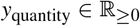 represent the profile and quantity predictions, respectively. We then calculated the loss as

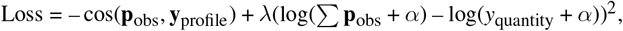

where *α* = 10^−6^ was used as a pseudocount and *λ* = 1/500 was used as a balancing weight between the profile and quantity loss functions. The models were fit using the Adam optimizer (learning rate = 0.001) with early stopping (patience = 10 epochs).

### Variant masking / reference sequence training

We retrained a CLIPNET model using the full set of 67 PRO-cap libraries, but instead of using the corresponding personalized genome sequences, we extracted sequences from the hg38 reference genome. The model architecture, hyperparameters, and data splits were as described above and in the CLIPNET paper [17].

### LCL benchmarks

We benchmarked the subsampled and reference-trained CLIPNET models using the tasks described in our previous manuscript [17].

Specifically, for the cross-loci benchmark, we considered a set of 4598 high confidence PRO-cap peaks on the holdout chromosomes. We extracted the hg38 reference sequence and the average RPM-normalized PRO-cap coverage around each of these peaks [19], then measured the performance of the models by the median Pearson’s correlation between predicted and observed PRO-cap coverage tracks (profile performance) and the Pearson’s correlation between the predicted and observed log_10_ total PRO-cap coverage (quantity performance). In our previous paper, we reported performance metrics for each model in the CLIPNET ensemble on its unique holdout set; for the sake of brevity here, we only report the performance of the ensemble on the chromosomes that were entirely withheld during training (chromosomes 9, 13, 20, and 21).

For the tiQTL and diQTL benchmarks, we used a set of 2,057 tiQTLs and 1,027 diQTLs that were previously fine-mapped by [19] and used to benchmark CLIPNET [17]. We measured the predicted and observed QTL effects by first binning individuals based on their QTL genotype, then taking the *L*^2^ norm of the difference vector between averaged homozygous reference and averaged homozygous alternative tracks. To avoid data leakage, we used the same QTL score compositing scheme we previously described [17]. Specifically, for QTLs on the completely withheld data fold 0, we used the predictions from the CLIPNET ensemble. For the QTLs on the remaining chromosomes, we used the prediction from the model replicate where that QTL was part of the hold out data fold.

### K562 fine tuning

We used fine tuning to adapt CLIPNET to a K562 PRO-cap dataset. We downloaded bigWig files (ENCSR261KBX) containing from the ENCODE data portal [36], then merged isogenic replicates. To maintain consistency across the PRO-cap datasets used in this study, we re-called PINTS peaks [32] rather than using the ENCODE peaks and applied an RPM transform to the PRO-cap tracks for training and evaluation. As this dataset was mapped to hg38 and K562 is a cancer cell line with a poorly defined karyotype, hg38 was used as the source of genome sequences for fine tuning.

The K562 models were initialized using the weights from the full personalized LCL model (keeping the same architecture), then trained to predict PRO-cap profiles and total coverage at K562 PINTS peaks. We opted to use fine tuning (i.e., all weights were set as trainable) rather than final layer probing, as CLIPNET is a relatively compact model. We used the same hyperparameters as when training the LCL models, but with a single initial warmup epoch at a learning rate of 0.0001. Additionally, to prevent potential data leakage in the variant effect prediction tasks, we used the same data splits as for the LCL model.

To test the contribution of pre-training on personalized genomic sequences, we also applied the same fine tuning procedures to the reference-trained CLIPNET LCL model.

### K562 PRO-cap prediction benchmark

We compared the performance of CLIPNET and ProCapNet at cross-loci prediction in K562. Since the two models used different holdout datasets, we decided to evaluate the prediction accuracies of individual CLIPNET and ProCapNet model folds on their respective holdout chromosomes. We calculated the median Pearson’s correlation between the predicted and observed profiles, as well as the Pearson’s correlation between the predicted and observed log_1_ 0 transcription initiation quantities (RPM-scaled for CLIPNET, raw counts for ProCapNet). The custom K562 peak call set used to train CLIPNET was used for both models. CLIPNET was evaluated on 1000 bp sequence windows and 500 bp tracks, whereas ProCapNet was evaluated on 2114 bp sequence windows and 1000 bp tracks.

### K562 MPRA benchmark

Siraj et al. recently characterized the impact of hundreds of thousands of non-coding variants in MPRAs across several human cell lines [23]. We chose to analyze the K562 MPRA dataset, as K562 is the only cell line tested in that study with published PRO-cap data, and which was used to train Enformer [9] and ProCapNet [10] models.

To test the ability of CLIPNET K562 (both personal and reference-pre-trained) to SNP effects in this MPRA, we reconstructed the MPRA sequences by inserting the tested 200 bp oligonucleotides (Supplemental Table 2 of [23]) directly upstream of the minimal TATA promoter of pMPRAv3:minP-GFP, (Addgene #109035) as described in [23]. We then extracted 1 kb of sequence from each reporter containing the oligonucleotide insert and the minimal promoter. We predicted the transcription initiation quantity for each allele and measured the predicted effect of each SNP as the log_2_ ratio of the reference and alternate allele predictions.

To generate ProCapNet [10] predictions, we simply padded the sequences constructed for CLIPNET out to the 2114 bp context length used by that model using the reporter backbone. We calculated the average predicted initiation quantity across the 7 ProCapNet K562 model replicates (ENCFF976FHE) for each reporter construct, then calculated the SNP effects as described for CLIPNET.

For the Enformer [9] predictions, we used the endogenous genomic sequences surrounding each SNP, as the long context length used for this model makes it impractical to test on the reporter sequences. We extracted 196,608 bp of sequence centered on each SNP (excluding those that would cause spillovers off the edge of the chromosome), then predicted K562 DNase (tracks 33, 34, 35, 121, 122, 123, and 625) and CAGE (tracks 4828, 5111) tracks in the central 8 bins (representing 1024 bp) using the PyTorch implementation of Enformer (https://github.com/lucidrains/enformer-pytorch/). After some testing, we determined that K562 DNase (specifically, track 122, corresponding to

ENCODE accession ENCFF868NHV) was the best predictor for MPRA SNP effect (as described below). We thus set the Enformer SNP predictions as the log_2_ ratio between the reference and alternative allele DNase predictions.

We considered the following two benchmarks for MPRA prediction. First, we measured the ability of each model to distinguish between expression modulating variants (emVar) and active non-emVars in K562 (Supplemental Table 3 of [23]). Specifically, we calculated precision-recall curves (PRC), using the square of the SNP prediction for each model as the predictor. To avoid possible data leakage from variants in the pre-training of the personalized CLIPNET model, we only considered variants on the holdout chromosomes for CLIPNET (chromosomes 9, 13, 20, and 21). We also excluded variants that would cause chromosome edge spillovers for Enformer, resulting in 2038 emVars and 4057 non-emVars. We observed that the personalized CLIPNET K562 model achieved the highest area under the PRC (auPRC). To measure the accuracy of this estimate, we performed *n* = 1000 bootstraps of the holdout SNPs, then calculated the difference in auPRC between all other models and the personalized CLIPNET K562 model.

Since both CLIPNET K562 models and ProCapNet K562 produce predictions of transcription initiation, we also quantitatively assessed the SNP effect predictions made by all three models. We generated scatterplots of the log_2_FC predictions made by each model against the observed log_2_FC values (specifically, mean RNA counts/mean DNA counts per allele). We also calculated Pearson’s correlation coefficients and sign mismatches between the predicted and observed log_2_FC values.

## Declarations

### Data availability

This study makes use of publicly available datasets. URLs and accession codes are provided in the methods section. The original CLIPNET model and the transfer-learned K562 model are available at https://doi.org/10.5281/zenodo.10408623 and https://doi.org/10.5281/zenodo.11196189, respectively. All other supporting data for this study, including model weights for the ablation tests, are available at https://doi.org/10.5281/zenodo.13823014.

### Code availability

Code for performing and plotting the results of the two ablation tests is available at https://github.com/Danko-Lab/clipnet_ablation. Code for fine tuning CLIPNET to predicting initiation in K562 and evaluating on the MPRA dataset are is available at https://github.com/Danko-Lab/clipnet_k562. This study also makes use of utility scripts developed for the original CLIPNET paper (https://github.com/Danko-Lab/clipnet).

### Author contributions

A.Y.H and C.G.D conceived of the project. A.Y.H designed the analyses with input from C.G.D. A.Y.H. and N.P.P. implemented the analyses. A.Y.H. and C.G.D. wrote the manuscript together.

## Acknowledgments

We thank Li Yao and Haiyuan Yu for providing a pre-publication version of their PINTS peak-calling software, Hojoong Kwak for assistance with interpreting his group’s QTL analysis, Kelly Cochran for advice on working with ProCapNet, and Anshul Kundaje for suggesting the variant masking ablation test.

## Supplementary Figures

**Figure S1:**
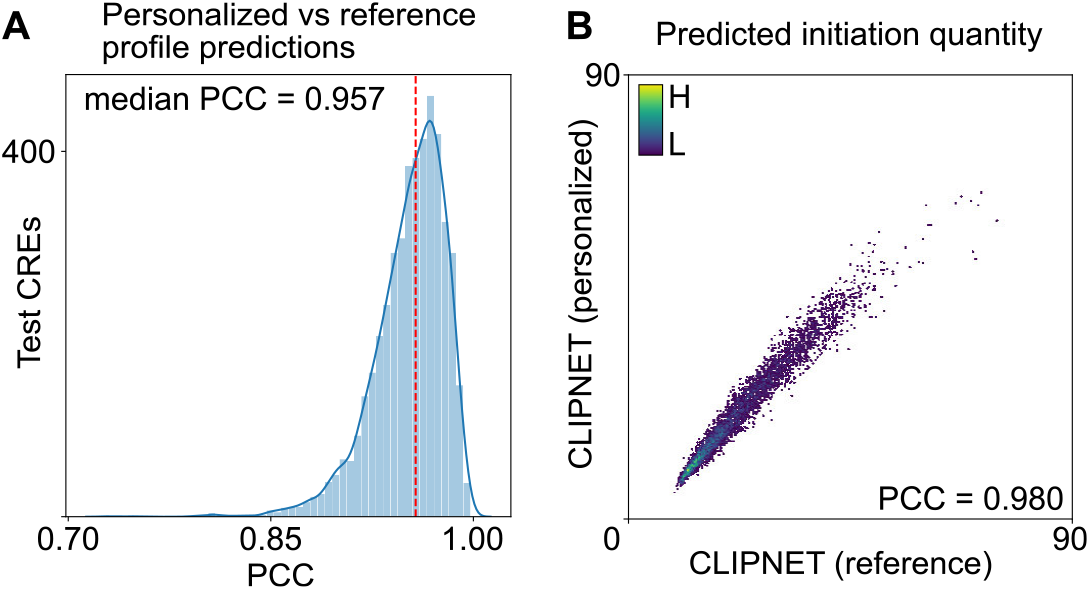
Comparison of reference-trained and personalized CLIPNET predictions in LCLs across genomic loci. (**A**) Initiation profiles predictions between reference-trained and personalized CLIPNET models are highly correlated (median profile Pearson’s correlation between predictions = 0.957). (**B**) Initiation quantity predictions between reference-trained and personalized CLIPNET models are highly correlated (Pearson’s correlation = 0.980). Points are colored by a Gaussian kernel density estimate.

**Figure S2:**
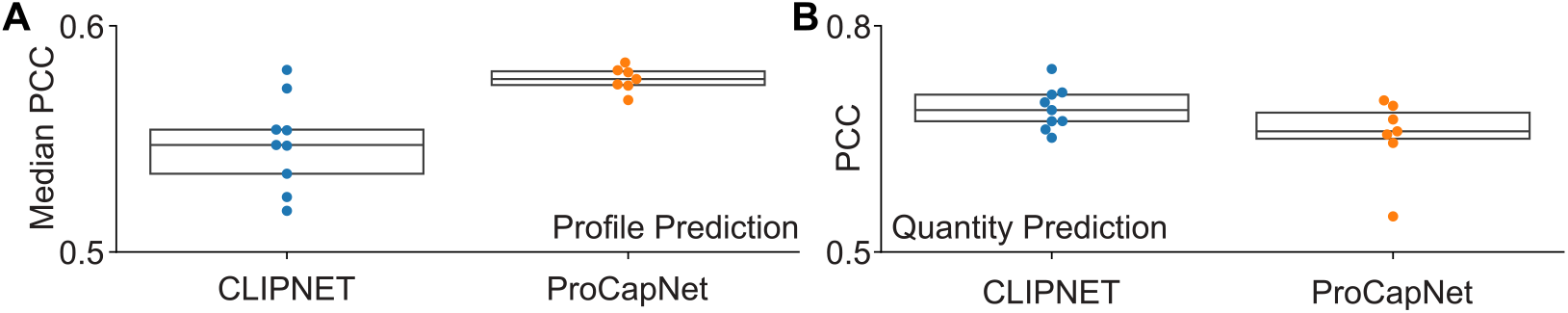
Comparison of CLIPNET and ProCapNet cross-loci predictions in K562. (**A**) Initiation profile prediction accuracy (median Pearson’s correlation of model replicates). (**B**) Initiation quantity prediction (Pearson’s correlation across model replicates). CLIPNET (*n* = 9) and ProCapNet (*n* = 7) models were evaluated on their respective holdout chromosomes. Box plots represent median and upper/lower quartiles.

## References

[1] Huang, C. et al. Personal transcriptome variation is poorly explained by current genomic deep learning models. Nat. Genet. 55, 2056–2059 (2023). URL http://dx.doi.org/10.1038/s41588-023-01574-w.

[2] Sasse, A. et al. Benchmarking of deep neural networks for predicting personal gene expression from DNA sequence highlights shortcomings. Nat. Genet. 55, 2060–2064 (2023). URL http://dx.doi.org/10.1038/s41588-023-01524-6.

[3] Karollus, A., Mauermeier, T. & Gagneur, J. Current sequence-based models capture gene expression determinants in promoters but mostly ignore distal enhancers. Genome Biol. 24, 56 (2023). URL http://dx.doi.org/10.1186/s13059-023-02899-9.

[4] Linder, J., Srivastava, D., Yuan, H., Agarwal, V. & Kelley, D. R. Predicting RNA-seq coverage from DNA sequence as a unifying model of gene regulation. Nat. Genet. 1–13 (2025). URL https://www.nature.com/articles/s41588-024-02053-6.

[5] Martyn, G. E. et al. Rewriting regulatory DNA to dissect and reprogram gene expression. bioRxiv 2023.12.20.572268 (2023). URL https://www.biorxiv.org/content/10.1101/2023.12.20.572268v1.

[6] Chandra, N. A., Hu, Y., Buenrostro, J. D., Mostafavi, S. & Sasse, A. Refining the cis-regulatory grammar learned by sequence-to-activity models by increasing model resolution. bioRxiv 2025.01.24.634804 (2025). URL https://www.biorxiv.org/content/10.1101/2025.01.24.634804v1.abstract.

[7] Pampari, A. et al. ChromBPNet: bias factorized, base-resolution deep learning models of chromatin accessibility reveal cis-regulatory sequence syntax, transcription factor footprints and regulatory variants. bioRxiv 2024.12.25.630221 (2024). URL https://www.biorxiv.org/content/10.1101/2024.12.25.630221v2.abstract.

[8] Avsec, Z. et al. Base-resolution models of transcription-factor binding reveal soft motif syntax. Nat. Genet. 53, 354–366 (2021). URL https://www.nature.com/articles/s41588-021-00782-6.

[9] Avsec, Z. et al. Effective gene expression prediction from sequence by integrating long-range interactions. Nat. Methods 18, 1196–1203 (2021). URL https://www.nature.com/articles/s41592-021-01252-x.

[10] Cochran, K. et al. Dissecting the cis-regulatory syntax of transcription initiation with deep learning. bioRxiv 2024.05.28.596138 (2024). URL https://www.biorxiv.org/content/10.1101/2024.05.28.596138v1.

[11] Dudnyk, K., Cai, D., Shi, C., Xu, J. & Zhou, J. Sequence basis of transcription initiation in the human genome. Science 384, eadj0116 (2024). URL https://www.science.org/doi/abs/10.1126/science.adj0116. https://www.science.org/doi/pdf/10.1126/science.adj0116.

[12] Zhou, J. et al. Deep learning sequence-based ab initio prediction of variant effects on expression and disease risk. Nat. Genet. 50, 1171–1179 (2018). URL https://www.ncbi.nlm.nih.gov/pmc/articles/PMC6094955/.

[13] Zhou, J. Sequence-based modeling of three-dimensional genome architecture from kilobase to chromosome scale. Nat. Genet. 54, 725–734 (2022). URL http://dx.doi.org/10.1038/s41588-022-01065-4.

[14] Bajwa, A., Rastogi, R., Kathail, P., Shuai, R. W. & Ioannidis, N. M. Characterizing uncertainty in predictions of genomic sequence-to-activity models. bioRxiv 279–297 (2023). URL https://proceedings.mlr.press/v240/bajwa24a.html.

[15] Drusinsky, S., Whalen, S. & Pollard, K. S. Deep-learning prediction of gene expression from personal genomes. bioRxiv 2024.07.27.605449 (2024). URL https://www.biorxiv.org/content/10.1101/2024.07.27.605449v1.abstract.

[16] Rastogi, R., Reddy, A. J., Chung, R. & Ioannidis, N. M. Fine-tuning sequence-to-expression models on personal genome and transcriptome data. bioRxiv 2024.09.23.614632 (2024). URL https://www.biorxiv.org/content/10.1101/2024.09.23.614632v1.abstract.

[17] He, A. Y. & Danko, C. G. Dissection of core promoter syntax through single nucleotide resolution modeling of transcription initiation. bioRxiv 2024.03.13.583868 (2024). URL https://www.biorxiv.org/content/10.1101/2024.03.13.583868.

[18] Core, L. J. et al. Analysis of nascent RNA identifies a unified architecture of initiation regions at mammalian promoters and enhancers. Nat. Genet. 46, 1311–1320 (2014). URL https://www.nature.com/articles/ng.3142.

[19] Kristjánsdóttir, K., Dziubek, A., Kang, H. M. & Kwak, H. Population-scale study of eRNA transcription reveals bipartite functional enhancer architecture. Nat. Commun. 11, 5963 (2020). URL http://dx.doi.org/10.1038/s41467-020-19829-z.

[20] Andersson, R. et al. An atlas of active enhancers across human cell types and tissues. Nature 507, 455–461 (2014). URL http://dx.doi.org/10.1038/nature12787.

[21] Tippens, N. D. et al. Transcription imparts architecture, function and logic to enhancer units. Nat. Genet. 52, 1067–1075 (2020). URL http://dx.doi.org/10.1038/s41588-020-0686-2.

[22] Lis, J. ENCSR261KBX (2021). URL https://www.encodeproject.org/experiments/ENCSR261KBX/. Title of the publication associated with this dataset: ENCODE Datasets.

[23] Siraj, L. et al. Functional dissection of complex and molecular trait variants at single nucleotide resolution. bioRxiv 2024.05.05.592437 (2024). URL https://www.biorxiv.org/content/10.1101/2024.05.05.592437v1.

[24] Abell, N. S. et al. Multiple causal variants underlie genetic associations in humans. Science 375, 1247–1254 (2022). URL http://dx.doi.org/10.1126/science.abj5117.

[25] Ray, J. P. et al. Prioritizing disease and trait causal variants at the TNFAIP3 locus using functional and genomic features. Nat. Commun. 11, 1237 (2020). URL https://www.nature.com/articles/s41467-020-15022-4.

[26] Tewhey, R. et al. Direct identification of hundreds of expression-modulating variants using a multiplexed reporter assay. Cell 165, 1519–1529 (2016). URL http://dx.doi.org/10.1016/j.cell.2016.04.027.

[27] Vockley, C. M. et al. Massively parallel quantification of the regulatory effects of noncoding genetic variation in a human cohort. Genome Res. 25, 1206–1214 (2015). URL http://genome.cshlp.org/content/25/8/1206.abstract.

[28] de Almeida, B. P., Reiter, F., Pagani, M. & Stark, A. DeepSTARR predicts enhancer activity from DNA sequence and enables the de novo design of synthetic enhancers. Nat. Genet. 54, 613–624 (2022). URL http://dx.doi.org/10.1038/s41588-022-01048-5.

[29] de Almeida, B. P. et al. Targeted design of synthetic enhancers for selected tissues in the drosophila embryo. Nature 626, 207–211 (2024). URL https://www.nature.com/articles/s41586-023-06905-9.

[30] Taskiran, I. I. et al. Cell-type-directed design of synthetic enhancers. Nature 626, 212–220 (2024). URL https://www.nature.com/articles/s41586-023-06936-2.

[31] Gosai, S. J. et al. Machine-guided design of cell-type-targeting cis-regulatory elements. Nature 634, 1211–1220 (2024). URL https://www.nature.com/articles/s41586-024-08070-z.

[32] Yao, L. et al. A comparison of experimental assays and analytical methods for genome-wide identification of active enhancers. Nat. Biotechnol. 40, 1056–1065 (2022). URL http://dx.doi.org/10.1038/s41587-022-01211-7.

[33] Tome, J. M., Tippens, N. D. & Lis, J. T. Single-molecule nascent RNA sequencing identifies regulatory domain architecture at promoters and enhancers. Nat. Genet. 50, 1533–1541 (2018). URL http://dx.doi.org/10.1038/s41588-018-0234-5.

[34] Lin, J., Luo, R. & Pinello, L. EPInformer: a scalable deep learning framework for gene expression prediction by integrating promoter-enhancer sequences with multimodal epigenomic data. bioRxiv 2024.08.01.606099 (2024). URL https://www.biorxiv.org/content/10.1101/2024.08.01.606099v1.abstract.

[35] Gasperini, M. et al. A genome-wide framework for mapping gene regulation via cellular genetic screens. Cell 176, 377–390.e19 (2019). URL https://pubmed.ncbi.nlm.nih.gov/30612741/.

[36] Luo, Y. et al. New developments on the encyclopedia of DNA elements (ENCODE) data portal. Nucleic Acids Res. 48, D882–D889 (2020). URL http://dx.doi.org/10.1093/nar/gkz1062.

